# Arctic plant species display contrasting levels of chloroplast DNA copy numbers

**DOI:** 10.1101/2025.01.29.635447

**Authors:** Stefaniya Kamenova, Sylvain Moinard, Samantha Paige Huset Dwinnell, Frédéric Laporte, Éric Coissac

## Abstract

DNA metabarcoding has revolutionised our ability to characterise biodiversity at unprecedented spatial and temporal scales, outperforming most traditional methods for biodiversity monitoring. However, DNA metabarcoding is not without limitations, in particular regarding the quantitative relationship between sequence read abundances and species numerical abundances or biomass (i.e., quantitative performance). Variation in DNA copy numbers has been often pinpointed as a potentially important factor biasing DNA metabarcoding quantitative performance but empirical comparisons of DNA copy number variation across species remain rare. Here, we identify chloroplast DNA copy number variation (cpDNA CNV) in plants as a potentially important factor that could impact quantitative performance in DNA metabarcoding studies. Using digital droplet PCR, we quantified chloroplast copy number variation in four common high Arctic plant species and between two plant tissue types (green tissues and roots). The amount of cpDNA per unit of dry mass varied by a factor of 3 (for roots) to 7.6 (for green tissues) among species, and up to 67 when comparing cpDNA copy numbers between green and root tissues from the same species. Despite significant differences in cpDNA copy numbers among species for both green tissues and roots, the most pronounced differences in cpDNA copy numbers were clearly between the two tissue types tested. These findings suggest that variation in cpDNA copy numbers among plant species and particularly plant tissue types can be an important but underestimated factor impacting plant sequence reads abundance in DNA metabarcoding datasets. We call for more extensive cpDNA CNV referencing efforts from wild-ranging plants to improve the use of DNA metabarcoding for research and biodiversity monitoring.

## Introduction

DNA metabarcoding has emerged as an indispensable tool for the study and monitoring of biodiversity (Valentini et al. 2009; Taberlet et al. 2012; Deiner et al. 2017; Ficetola & Taberlet 2023). The method has been applied in a variety of contexts, from biodiversity surveys to diet analyses to the reconstruction of ancient ecosystems (Ando et al. 2020; Ariza et al. 2022; Deagle et al. 2017; de Sousa et al. 2019; Edwards et al. 2018; Foote et al. 2012; Kjær et al. 2022; Leray & Knowlton 2015; Thomsen et al. 2012). The success of DNA metabarcoding is mainly driven by its high versatility as well as cost- and time-effectiveness. High detection sensitivity adds up to the list of advantages, often making DNA metabarcoding the only option for tracking rare or highly elusive species (Jerde et al. 2011; Fediajevaite et al. 2021; Furlan et al. 2019; Parker et al. 2022; Pochon et al. 2013).

As every method, DNA metabarcoding has limitations, namely quantitative performance – i.e., the extent to which sequence reads abundances correlate with species numerical abundances or biomass. DNA metabarcoding can be quantitative (Lamb et al. 2019) but there is also substantial variation among case studies, highlighting the lack of understanding regarding the underlying factors. DNA metabarcoding involves a relatively large number of analytical steps such as sampling, DNA extraction, PCR, sequencing, and bioinformatic analyses — each contributing to inject errors into DNA metabarcoding datasets. Several studies have already pinpointed issues related to differences in detection probabilities among species, PCR stochasticity as well as biases in the analytical pipelines (Yang et al. 2021; Luo et al. 2022; Moinard et al. 2023). Efforts to account for one or multiple of biases along the DNA metabarcoding pipeline can help (cf Moinard et al. 2023) but does not always lead to an improved quantitative information (Krehenwinkel et al. 2017; Yang et al. 2021). In fact, studies controlling for PCR stochasticity and primer-template mismatch biases suggest that discrepancy in DNA quantitative estimates might be driven by differences in the copy number variation (CNV) among species (Thomas et al. 2015; Krehenwinkel et al. 2017; Garrido-Sanz et al. 2020; 2022). This phenomenon has already been documented in prokaryotes for the SSU rRNA 16S gene, with copy numbers showing considerable variation among bacterial taxa (Farrelly et al. 1995; Fogel et al. 1999; Kembel et al. 2012; Angly et al. 2014). In a recent study, Vasselon et al. (2017) revealed significant differences in chloroplast *rbc*L gene copy numbers among eight species of benthic diatoms, also correlating with the differences in cell biovolume. Yet, formal assessments for multicellular organisms are still missing.

In plants, chloroplast genes are a preferred target for DNA metabarcoding (Taberlet et al. 2007; Valentini et al. 2009), especially for the analysis of herbivores diets (Soininen et al. 2009; Pansu et al. 2022; Spitzer et al. 2023; Ratkiewicz et al. 2024). Still, it remains unclear how potential variation in chloroplast DNA (cpDNA) copy number in plants might be a factor impacting the quantitative performance of DNA metabarcoding. Differences in cpDNA CNV among plant taxa have been well-documented (Bendich 1987; Li et al. 2006; Zoschke et al. 2007; Bennett & Leitch 2012), together with the understanding that cpDNA can vary dynamically with plant development and tissue type (Boffey & Leech 1982; Baumgartner et al. 1989; Sakamoto & Takami 2018; Domínguez & Cejudo 2021). Such variation is a function of the number of plastids per cell and the amount of cpDNA per plastid (Boffey & Leech 1982).

Here, using droplet digital PCR, we compare cpDNA CNV in four plant species, commonly occurring within the High Arctic tundra of the Svalbard archipelago. We focus on species with contrasting life-history traits: grass (*Poa arctica*), sedge (*Carex subspathacea*), shrub (*Salix polaris*) and one small dicotyledon species (*Saxifraga oppositofolia*). Specifically, we quantify cpDNA CNV in both roots and green tissues for all species. For *Saxifraga oppositofolia*, we also assessed cpDNA copy numbers of the flowers. We hypothesized significant differences in cpDNA CNV among the four species as well as significantly higher cpDNA copy numbers in green tissues compared to roots and flowers.

## Material & Methods

### Field sampling

Fieldwork took place in Nordenskiöld land in Svalbard (77°50’–78°20’N, 15°00’–17°30’E) during a period of one week in July 2022, coinciding with the plant peak greenness period in this area. All species and individuals were sampled in close vicinity to minimise the potential effects of spatial variation. For each species, we sampled whole individuals, including all the green tissues and the roots as well as flowers for *Saxifraga oppositofolia* (see sample size in Table S1). Plant individuals were dug out from the soil with the roots, soil particles removed, and each plant was immediately placed in an individual zip-lock plastic bag prefilled with silica gel beads (Carl Roth, Germany). Samples were shipped to University of Oslo for further analysis. Each plant specimen was carefully dislodged from the silica gel beads, weighed, and different tissue types (i.e., flowers, roots, and green tissues, including stems, leaves, or both) subsampled separately. Between 5 and 25 mg of silica-dried tissue was retrieved for DNA extraction for the different tissue types depending on the size and the biomass of the plant (Table S2). The remainder of plant specimens was oven-dried at 60°C for 48 hours and the remaining tissue mass was weighed again. For one sample, the low biomass did not allow additional weighting (so6f). The ratio of fresh-to-dry weight for these two samples was therefore estimated by using the median of the fresh-to-dry weight ratio among all samples.

### Quantifying cpDNA CNV with digital droplet PCR

Plant tissue samples were homogenised using liquid nitrogen and sterile mortar and pestle. DNA was extracted using the NucleoSpin Plant II (Macherey-Nagel, Germany) and DNA concentrations assessed with NanoDrop 1000 spectrophotometer (Thermo Fisher Scientific, Massachusetts, USA), and subsequent serial dilutions of DNA were carried out based on these values. Plant DNA was amplified using the *Sper01* primer set (Bonin et al. 2018) and the copies of the trn*L* P6 loop g-h DNA fragment (Taberlet et al. 2006) were quantified by droplet digital PCR (ddPCR) using the QX200 Droplet Digital™ System (Bio-Rad). Thermocycler conditions with optimised annealing temperature for the *Sper01* primers were set according to the QX200 ddPCR EvaGreen Supermix user manual for 3-step PCRs. The final program started with 5 min of activation at 95 °C, followed by 40 cycles of 30 s denaturation at 95 °C, 1 min annealing at 58 °C and 1 min extension at 72 °C, followed by a first step of stabilisation at 4 °C for 5 min and a second step at 90 °C for 5 min and finished with 12 °C on hold. The analysis was performed using the Absolute Quantification (ABS) mode in a final volume of 20 μl of reaction mix. This allows measurement of target DNA concentrations ranging from less than 0.25 copies/μl to more than 5 000 copies/μl (Droplet Digital PCR Applications Guide, Biorad). The dilution range was performed for each sample for 0.25 ng/μl, 0.025 ng/μl and 0.0025 ng/μl of total DNA. Two replicates were done for each dilution level. According to the documentation, the assay is not accurate above several thousand copies/μl. In the cases where the intermediate dilution (0.025 ng/μl of sample) was above 750 copies/μl, the concentration for the most concentrated level was expected to be above 7 500 copies/μl, and therefore outside the measurement range. Ten replicates were discarded based on this criterion. Three replicates with an abnormally low signal were also discarded. Details about all the measures retained for further analyses are presented in Table S3.

### Data analyses

Data analyses were carried out with R (version 4.4.1). Correction factor for sequence reads abundance was estimated by converting the measured target DNA concentrations to a number of target copies per mg of dry weight. This is calculated for each replicate using the equation 1:

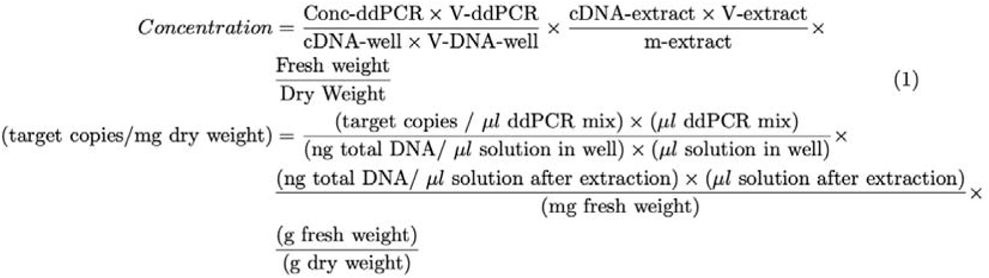

where “Conc-ddPCR” corresponds to the concentration (copies/μl) assayed by ddPCR, “V-ddPCR” to the total volume of ddPCR reaction mix (20 μl), “V-DNA-well” to the volume of DNA solution in the ddPCR reaction mix (5 μl), “cDNA-well” to each of the three dilutions (0.25, 0.025 or 0.0025 ng/μl), “cDNA-extract” to the DNA concentration after DNA extraction (in ng/μl), “V-extract” to the total volume of extracted DNA solution (100 μl), “m-extract” to the fresh weight of plant used for the extraction (in mg), “Fresh Weight” to the grams of fresh plant biomass before oven-drying, and finally “Dry Weight” corresponding to the grams of dry plant biomass after oven-drying. Median concentrations were compared among species for a given tissue and among tissues for a given species using Wilcoxon tests for 2-groups comparisons and Kruskal-Wallis tests (reported with H-statistics and degrees of freedom), followed whenever necessary by *post hoc* Dunn tests for multiple comparisons. The p-values were corrected by Bonferroni’s method for multiple comparisons and compared to the 0.05 threshold.

## Results

The median dry-to-fresh mass ratio was 0.954 for all samples and was not significantly different between tissue types (0.947 for flowers, 0.954 for roots and 0.953 for green tissues, p-value = 0.56). We observed in average 27 times more cpDNA copy numbers in green tissues compared to roots (p-value < 0.001) (Table 1, Figure 1), with, at maximum, *Carex subspathacea* having cpDNA copy number values 67 times higher in green tissues compared to roots (p-value < 0.001). cpDNA copy numbers in green tissues were 14 times higher compared to cpDNA copy numbers in roots for *Saxifraga oppositofolia* (p-value < 0.001), 19 times higher for *Salix polaris* (p-value < 0.001) and 6.2 times higher for *Poa arctica* (p-value < 0.001). Moreover, in *Saxifraga oppositofolia*, cpDNA copy numbers did not differ between roots and flowers (p-value = 1), but cpDNA copy numbers in green tissues were in median 16 times higher compared to flowers (p-value < 0.001). There were significant differences in cpDNA copy numbers in green tissues among plant species (p-value < 0.001, H = 37.6 with 3 degrees of freedom), with the highest difference being between *Carex subspathacea* and *Saxifraga oppositofolia*, the former having 7.6 times more cpDNA copy numbers per unit of dry mass (p-value < 0.001). Overall, cpDNA copy numbers in root tissues did differ among plant species (p-value = 0.003, H = 13.9 with 3 degrees of freedom) with, at maximum, cpDNA copy numbers being 3.0 times higher in *Poa arctica* roots compared to *Saxifraga oppositofolia* (p-value = 0.02) and 2.0 higher in *Poa arctica* roots compared to *Carex subspathacea* roots (p-value = 0.011). The other pairwise comparisons for roots were not significant.

**Table 1.**
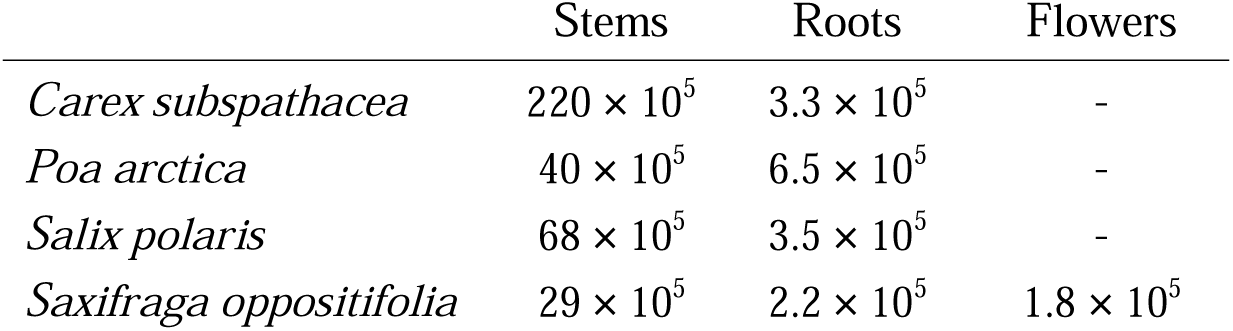
Median chloroplast DNA copy numbers per unit of dry mass across species and plant tissues in the four high Arctic plant species analysed.

**Figure 1.**
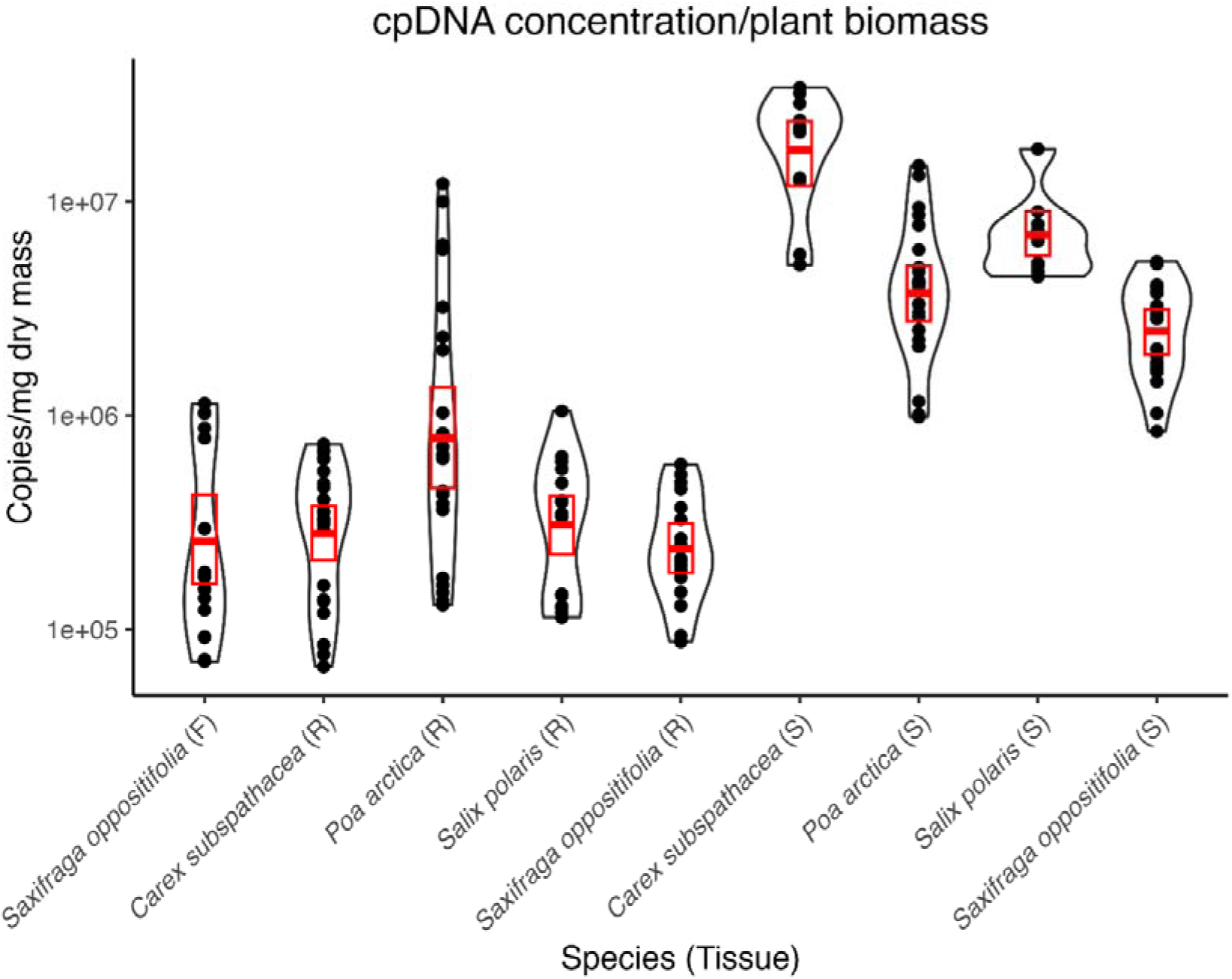
Variation in chloroplast copy numbers per unit of dry mass across species and plant tissue types in the four high Arctic plant species analysed. F=flowers, R=roots, S=green tissues.

## Discussion

We quantified chloroplast DNA copy number variation (cpDNA CNV) in four common Arctic plant species sampled during the peak greenness period. As expected, results show significant variation in cpDNA copy numbers, both among plant species and plant tissue types. There were up to 7.6-fold differences in cpDNA copy numbers per unit of dry mass in green tissues among the four species, and up to 67-fold differences between green tissues and roots within the same species. *Salix polaris* and *Carex subspathacea* displayed the highest concentrations in cpDNA for green tissues, while *Poa arctica* displayed the highest cpDNA concentrations in roots. Compared to green tissues, significantly less cpDNA copy numbers per unit of dry mass were detected in roots for all plant species as well as in flowers for *Saxifraga oppositofolia*.

cpDNA CNV in plant cells is known to be affected by three main factors: (i) the number of chloroplasts per cell; (ii) the number of genomes per chloroplast; and (iii) the number of copies per chloroplast genome of the target gene marker. Chloroplasts are concentrated in the mesophyll tissue and the size and the number of chloroplasts and genome copies per chloroplast gradually increase with tissue maturation (Boffey & Leech, 1982; Baumgartner et al.1989; Rauwolf et al. 2010; Greiner et al. 2019). Typically, cpDNA copy numbers are the lowest in juvenile meristematic cells (75–120 copies) and quickly stabilize in mature mesophyll tissues (2750–3200 copies) (Greiner et al. 2019). Interestingly, these values seem to be remarkably similar across species as different as sugar beet (*Beta vulgaris*, Amaranthaceae), arabidopsis (*Arabidopsis thaliana*, Brassicaceae), tobacco (*Nicotiana tabacum*, Solanaceae) or maize (*Zea mays*, Poaceae) (Greiner et al. 2019 and references therein). However, other studies report that DNA amounts for fully developed chloroplasts can span almost three orders of magnitude, from several hundred in species such as *A. thaliana* and wheat (*Triticum aestivum*, Poaceae) to more than ten thousand copies in wheat, or barley (*Hordeum vulgare,* Poaceae) (Boffey & Leech 1982; Leutwiler et al. 1984; Zoschke et al. 2007; Liere & Börner 2013). This discrepancy in cpDNA CNV estimates across different studies perhaps reflects at least partly differences in methodology (cf Bowman 1986; Li et al. 2006).

The chloroplast size and numbers also depend on the balance between plant cells growth and division as well as chloroplast division and cpDNA synthesis (Rauwolf et al. 2010), depending on plant metabolic needs and phenological stage at the moment of sampling. We tried to minimise variation by sampling plants in close vicinity to each other and within a period of few days. But even within the same experimental conditions, high Arctic plants do display differences in the rate and onset of senescence (Iveland 2023), with impact on cell growth and chloroplast division, which could explain the differences in cpDNA CNV among the four species here. Moreover, differences in moisture levels and nutrients availability are important factors modulating the rate of senescence through differences in nitrogen availability, also underpinning chlorophyll content and photosynthetic capacity (Semenchuk et al. 2015). On Svalbard, Poaceae are typically associated with mesic habitats, while *Carex subspatacaea* is common in marshes, and *Salix polaris* occurs in the moisture-rich *Luzula* heath and wet moss habitats (Dwinnell et al. 2024), whose high humidity and nutrients accumulation could explain the higher cpDNA CNV observed here for these two species.

Another important aspect to consider is the fact that we have analysed bulk leaf and stem parts, representing a complex mixture of mesophyll and non-mesophyll tissues (e.g., epidermis, vascular cells), with the latter containing less and smaller chloroplasts, typically carrying significantly less cpDNA copies (Rowan et al., 2009; Liere & Börner 2013). It is possible that differences in cpDNA CNV observed here might be jointly driven by differences in senescence rates or habitat use, and differences in the mesophyll to non-mesophyll tissues ratio at the moment of sampling, potentially contributing to increase inter- and intra-species variation in cpDNA. The ratio between leaves and stems that could be sampled for DNA extraction was difficult to standardise due to the small size of the plants as well as the limited biomass per individual, contributing to explain the intraspecific variability in cpDNA CNV here. To address these aspects, similar analysis could be carried out using plants cultured in controlled conditions akin to the approach by Vasselon et al. (2017) with the freshwater diatoms.

The most marked differences in cpDNA CNV appear to be between the green and the root tissues as well as between green tissues and flowers for *Saxifraga oppositofolia*. We observed up to two orders of magnitude less cpDNA copy numbers in roots and flowers across the four plant species, suggesting these tissue types contain either less plastids and/or less DNA copy numbers per plastid. Chloroplasts are abundant in mature photosynthetic tissues, while storage-specialised plastids (amyloplasts) are dominate in the roots. Plastids from the same tissue can also dynamically interconvert in one type or another in response to developmental or environmental cues (Jarvis & López-Juez 2013; Liebers et al. 2017; Choi et al. 2021; Domínguez & Cejudo 2021). However, identical copies of the same genome are present in all differentiated plastid types (Li et al. 2006). Studies comparing plastid DNA CNV among plant tissues remain rare, but the lower plastid DNA CNV in roots compared to photosynthetic tissues has been confirmed for two other plant species - maize (*Zea mays*, Ma & Li 2015) and the sago palm (*Metroxylon sagu*, Lim et al. 2020), while we could not find any such comparisons for flowers.

Considering the limited research on the topic, assessing the ecological consequences of our findings seems difficult. Given the large differences in cpDNA copy numbers reported here, both among species and among tissue types, it is legitimate to think of cpDNA CNV as a potential source of bias with DNA metabarcoding. Specifically, we would expect that within the same sample (e.g., herbivore faeces, soil, herbal mixtures), species with high cpDNA copy numbers will have their relative reads abundance overestimated. This bias will be even stronger for species with high cpDNA CNV differences, if different tissue types are mixed within the same sample. Therefore, assessing the extent to which cpDNA CNV impacts DNA metabarcoding quantitative performance is indispensable. For instance, an important priority is to test whether cpDNA copy numbers correlate with sequence reads abundance. For example, Neidel & Traugott (2023) combined ddPCR and DNA metabarcoding to track the DNA decay of ingested common dandelion seeds from the gut contents of the carabid *Pseudoophonus rufipes*. Results show that over time, dandelion cpDNA copy numbers positively correlated with DNA metabarcoding sequence reads abundance. But this relationship has never been tested for meal mixtures including different species and tissue types. Moreover, wild-ranging herbivores feed on a much larger array of plant tissues at different phenological stages, also comprising different cell types and cell density, meaning the relationship between cpDNA CNV and sequence reads abundance could be more complex. Feeding trials using simple diets of two or three dietary items show either good (Willerslev et al. 2014), or poor (Neby et al. 2021; Stapleton et al. 2022) agreement between sequence read abundances and ingested biomass meaning more specific parameters regarding the experimental meal have to be considered. Quantifying the part of variation in DNA metabarcoding quantitative performance explained by differences in cpDNA CNV can help explaining the incongruity among studies so far. The topic is also relevant for sedimentary or soil DNA metabarcoding studies. Typically, in such studies the primary target is extra-cellular plant DNA (Pietramellara et al. 2009; Torti et al. 2015; Zinger et al. 2016; Giguet-Covex et al. 2019) but investigating whether sequence reads abundances from extra-cellular plant DNA are sensitive to initial differences in cpDNA copy numbers between species and tissue types preserved in the soil or the sediments could be equally important.

Overall, our findings here also raise the question about the predictability of cpDNA CNV in natural conditions, also conditioning potential future referencing efforts as well as the application of correction factors for DNA metabarcoding analysis. For example, it is unknown whether cpDNA CN values are species-specific and to what extent they vary across seasons and spatial gradients. Future assessments could also consider the relevance of possible alleviation strategies such as replacing chloroplast DNA markers with nuclear ribosomal RNA markers (e.g., ITS, Moorhouse-Gann et al. 2018; Prieto et al. 2024), even though ribosomal RNA genes in plants are also known to display variability in copy numbers (Rogers & Bendich 1987).

In conclusion, our proof-of-concept study shows significant differences in cpDNA copy numbers among plant species and tissues and pinpoints cpDNA CNV as a potentially important but still largely overlooked source of bias in DNA metabarcoding analysis. As our results open more questions than they allow to answer, we encourage further research efforts on the topic to both, enable better apprehension of the importance of cpDNA CNV as a source of bias and improve the robustness and interpretability of DNA metabarcoding data.

## Acknowledgements

We thank the Governor of Svalbard for permission to undertake the research as well as the technical staff at the University Centre in Svalbard (UNIS) and the Norwegian Polar Institute for support with the field campaign and access to facilities. This received support from the Research Council of Norway through projects 315454 (“PRISM: Understanding climate change impacts in an Arctic ecosystem: an integrated approach through the prism of Svalbard reindeer”) and 337271 (“PIECEMEAL: Cracking the diet of the Svalbard reindeer by integrating old and modern tools”), the Svalbard Environmental Fund (project 23R17050 “Building DNA density database for Svalbard flora”) as well as by the French National Funding Agency through project ANR-20-CE02-0020 (“ALPALGA: Assessing the biodiversity of algal microalgae and understanding their interaction with the environment”). SK was also supported by the Åsgard mobility programme, jointly funded by the French Institute and the Research Council of Norway.

## Data availability statement

All data are provided in the Supplementary material (Tables S1 to S3). Data analysis scripts are available at: https://github.com/LECA-MALBIO

## Author’s contributions

SK, SM, FL and ÉC conceived research. SPHD carried out plant sampling. FL and SK carried out molecular analyses. SM and ÉC analysed the data. SK wrote the manuscript with input from all authors.

